# Thematic Shifts in Early-High-Impact Cancer Genomics and Diagnostics Research: A Bibliometric and Semantic Analysis

**DOI:** 10.64898/2026.07.04.736459

**Authors:** Zheng Su, Tinsley Li

**Affiliations:** AnyHelix Ltd., The Cloud, Tai Kok Tsui, Hong Kong, China; Computational Biology Group, FAB Tech, Yantian, Shenzhen, Guangdong, 518083, China

## Abstract

Cancer genomics and diagnostics is a rapidly evolving field in which identifying which topics attract early citation prominence can inform laboratory investment, clinical translation, and research strategy. We developed a bibliometric framework to identify and characterize the most influential recent publications in this domain across two consecutive annual cohorts. Using a mathematically exact threshold-expansion algorithm, we ranked over 10,000 OpenAlex-indexed research articles per cohort by 18-month post-publication citation count. Large language model (LLM)-based topical relevance filtering yielded 50 substantively on-topic papers per cohort (100 total). LLM-based concept extraction and a two-stage, embedding-guided normalization pipeline produced 1,090 canonical concepts organized into 77 parent themes, enabling structured cross-cohort comparison of paper-level concept prevalence. The most cited papers in both cohorts were large-scale genomic infrastructure resources rather than single-disease mechanistic studies. Between consecutive cohorts, normalized frequencies increased most for clonal evolution and intratumoral heterogeneity, single-cell and spatial omics technologies, spatial transcriptomics, immune cell infiltration, and copy number variation, while liquid biopsy and ctDNA-related themes showed the largest declines. These findings indicate that early citation impact in cancer genomics is shifting toward integrative, spatially resolved, and heterogeneity-aware research, and demonstrate that LLM-augmented citation ranking provides a replicable, semantically enriched lens for monitoring thematic evolution in precision oncology. A web interface for exploring the results is available at https://pri.pepkio.com/.

## Introduction

Molecular characterization has become integral to contemporary cancer care. Tumour genomic profiling refines disease classification, identifies patients who may benefit from targeted or immunotherapeutic agents, and supports screening for germline variants associated with heritable cancer risk^1^. As sequencing costs have declined and therapeutic options have expanded, cancer genomics and diagnostics have emerged as a rapidly evolving interface between laboratory science, bioinformatics, and clinical decision-making. Yet the translation of genomic knowledge into routine practice remains uneven, shaped by biological complexity, implementation barriers, and persistent uncertainties about how best to match molecular findings to individual patients^2^. Understanding how this interdisciplinary field is prioritizing and integrating its constituent technologies is therefore relevant to researchers, diagnostic laboratories, health systems, and policy makers seeking to align infrastructure investment with emerging clinical needs.

The current diagnostic landscape spans multiple complementary modalities. Tissue-based next-generation sequencing (NGS) is widely used to guide management in advanced malignancies, with accumulating evidence that genotype-directed therapy can improve progression-free and overall survival compared with non-matched approaches^3^. More comprehensive whole-genome and transcriptome sequencing workflows are increasingly proposed as unified platforms capable of replacing cascade testing strategies and capturing a broader spectrum of clinically relevant aberrations^4^. Minimally invasive liquid biopsy approaches, particularly analysis of circulating tumour DNA (ctDNA), enable serial assessment of tumour genomes, treatment response, minimal residual disease, and emerging resistance without repeated tissue sampling^5,6^. Diagnostic paradigms are also extending beyond tumour-cell-centric molecular testing toward biomarkers derived from the tumour microenvironment (TME), reflecting growing recognition that stromal, immune, and vascular components influence prognosis and therapeutic response^7^. Artificial intelligence (AI) applied to digital pathology offers an additional route to infer molecular alterations and prognostic features from routine histopathology, potentially complementing or triaging molecular assays^8^. At the population level, large clinico-genomic registries such as the AACR Project GENIE harmonize real-world sequencing data across institutions, supporting trial matching, driver discovery in rare tumours, and evaluation of genome-guided care in diverse patient populations^9^. Together, these developments indicate that cancer genomics and diagnostics is not a single technology but a converging ecosystem of sequencing platforms, biomarker modalities, computational tools, and data resources.

The pace of innovation in this ecosystem carries both scientific and clinical implications. Despite guideline endorsement of molecular testing, substantial gaps persist along the precision oncology pathway—from tumour biopsy and biospecimen evaluation through biomarker ordering, result reporting, and treatment selection—with many potentially eligible patients failing to receive genotype-matched therapy^10^. Diagnostic laboratories and health systems must therefore anticipate which technologies, biomarker classes, and analytical frameworks are gaining prominence in the research literature, as these signals often precede broader clinical adoption. Bibliometric indicators, while imperfect proxies for scientific quality, provide one means of gauging which lines of inquiry attract near-term attention within the research community. In fast-moving fields such as cancer genomics, early citation visibility among highly cited papers may be especially informative, because foundational datasets, methods papers, and platform releases can accumulate citations rapidly and redirect downstream investigation across disease contexts.

Existing bibliometric studies have mapped publication growth, collaboration networks, and keyword trends in cancer omics and related domains, typically drawing on large corpora from the Web of Science or Scopus and applying co-occurrence, co-citation, or burst-detection analyses^11–13^. These corpus-wide approaches have documented shifts toward multi-omics integration, immunotherapy-oriented biomarkers, machine learning, and TME profiling. However, by weighting all publications within a domain equally and relying predominantly on author-assigned or indexer-derived keywords, such methods may not resolve which substantively on-topic papers exert the greatest near-term influence, nor how the semantic content of that influential subset evolves across consecutive publication years. Topic classification in open bibliographic indexes, including OpenAlex, further introduces imprecision: articles assigned to a given research topic may include work only tangentially related to the target domain^14^. To our knowledge, no existing framework combines exact ranking of papers by early post-publication citation impact, large language model (LLM)–based filtering for substantive topical relevance, and author-preserving concept normalization into a structured taxonomy amenable to longitudinal cohort comparison within Cancer Genomics and Diagnostics.

We hypothesized that the thematic composition of early-high-impact, substantively on-topic publications in this field shifts measurably between consecutive publication cohorts, and that such shifts can be captured through semantically enriched concept mapping rather than keyword co-occurrence alone. To test this, we developed a bibliometric framework centred on an 18-month Publication Rank Index that identifies top-cited articles within the OpenAlex topic “Cancer Genomics and Diagnostics” (topic identifier T11287) using a mathematically exact threshold-expansion algorithm. Candidate papers were evaluated for substantive topical relevance by LLM classification of titles and abstracts, and scientific concepts were extracted and normalized through a two-level taxonomy comprising canonical topics and higher-level parent themes, with embedding-guided batching to preserve author terminology while enabling cross-cohort comparison. Although LLMs can scale annotation tasks in scientometric research, their outputs represent structured approximations that benefit from external validation and should not be treated as ground truth^15^. This design prioritizes what is influential in the near term among on-topic work and addresses a complementary question to corpus-wide trend analyses: not what the field publishes most abundantly, but what the research community cites most prominently within a defined post-publication interval.

In this study, we aimed to identify and characterize the most influential recent research articles in Cancer Genomics and Diagnostics by 18-month citation impact; to apply LLM-based topical filtering and concept extraction to map the thematic landscape of top-ranked papers in two consecutive publication cohorts comprising articles published in 2023 and 2024; and to compare concept prevalence between cohorts to describe how research priorities among early-high-impact work may be evolving.

## Methods

### Study design and data source

This study applied a bibliometric framework to identify the most influential recent research articles in the OpenAlex topic “Cancer Genomics and Diagnostics” (topic identifier T11287) and to characterize the thematic landscape of early-high-impact publications across two consecutive publication cohorts. Bibliographic records were retrieved from OpenAlex^16,17^, an open scholarly database that indexes metadata, abstracts, citation counts, and topic classifications for published scientific works. Analyses were restricted to research articles classified under topic T11287, and two non-overlapping publication date ranges were queried: 1 January 2023 through 31 December 2023 for the Class of 2025 cohort, and 1 January 2024 through 31 December 2024 for the Class of 2026 cohort. Bibliographic data were retrieved from OpenAlex on 1 July 2026. These cohort labels reflect the calendar year in which each group’s 18-month post-publication citation window was completed. Review articles and other non-article document types were excluded from both cohorts at retrieval.

### Publication Rank Index and 18-month citation scoring

Scientific influence was quantified using an 18-month post-publication citation window^18,19^. For each candidate paper, the citation count was defined as the number of citing works whose publication dates fell within the 18-month interval beginning on the paper’s own publication date. Because bibliographic databases expose only citing-work publication dates rather than event timestamps of individual citations, this metric captures works published within the specified window rather than citations recorded during that interval. All papers in each cohort were required to have completed their full 18-month window before citation scoring, with window eligibility assessed relative to the 1 July 2026 data retrieval date.

To identify the top-ranked papers by 18-month citation count without scoring the entire field corpus, a threshold-expansion algorithm was employed^20^. The algorithm targets N on-topic papers for downstream analysis; the large language model (LLM) relevance filtering step (described below) requires a broader evaluation pool of 1.8N papers with scored 18-month citations, so the citation-scoring phase first targeted 90 papers per cohort (1.8 × 50). Within the citation-scoring step, an initial candidate set S was assembled by retrieving the top kN papers ranked by lifetime (total) citation count from OpenAlex, where k was set to 1.5 (yielding 135 papers as the initial citation-scoring set). Exact 18-month citation counts were then computed for all eligible papers in S by querying OpenAlex for citing works with publication dates within the 18-month window. The 18-month citation count of the N-th ranked paper in S established a threshold m, below which no paper with fewer total lifetime citations could enter the final leaderboard, because lifetime citation counts provide an upper bound on 18-month citations. An expanded candidate pool T was constructed by retrieving all papers with lifetime citation counts at or above m. For the Class of 2026 cohort, the 18-month threshold m was 45 citations, yielding an expanded pool of 191 papers; for the Class of 2025 cohort, m was 42, yielding an expanded pool of 351 papers. Exact 18-month citation counts were computed for all papers in T not already scored from S, and the final ranked list was derived by sorting the complete scored pool by 18-month citation count in descending order. This algorithm is mathematically exact and guarantees that no paper in the true top-N leaderboard is omitted.

### Topical relevance filtering

Because OpenAlex topic classification may include articles with only tangential connection to the target research domain, each candidate paper was evaluated for substantive topical relevance using a large language model (LLM)^21^. For each paper, the model was prompted with the paper’s title and abstract using the following classification instructions (with the topic set to Cancer Genomics and Diagnostics):

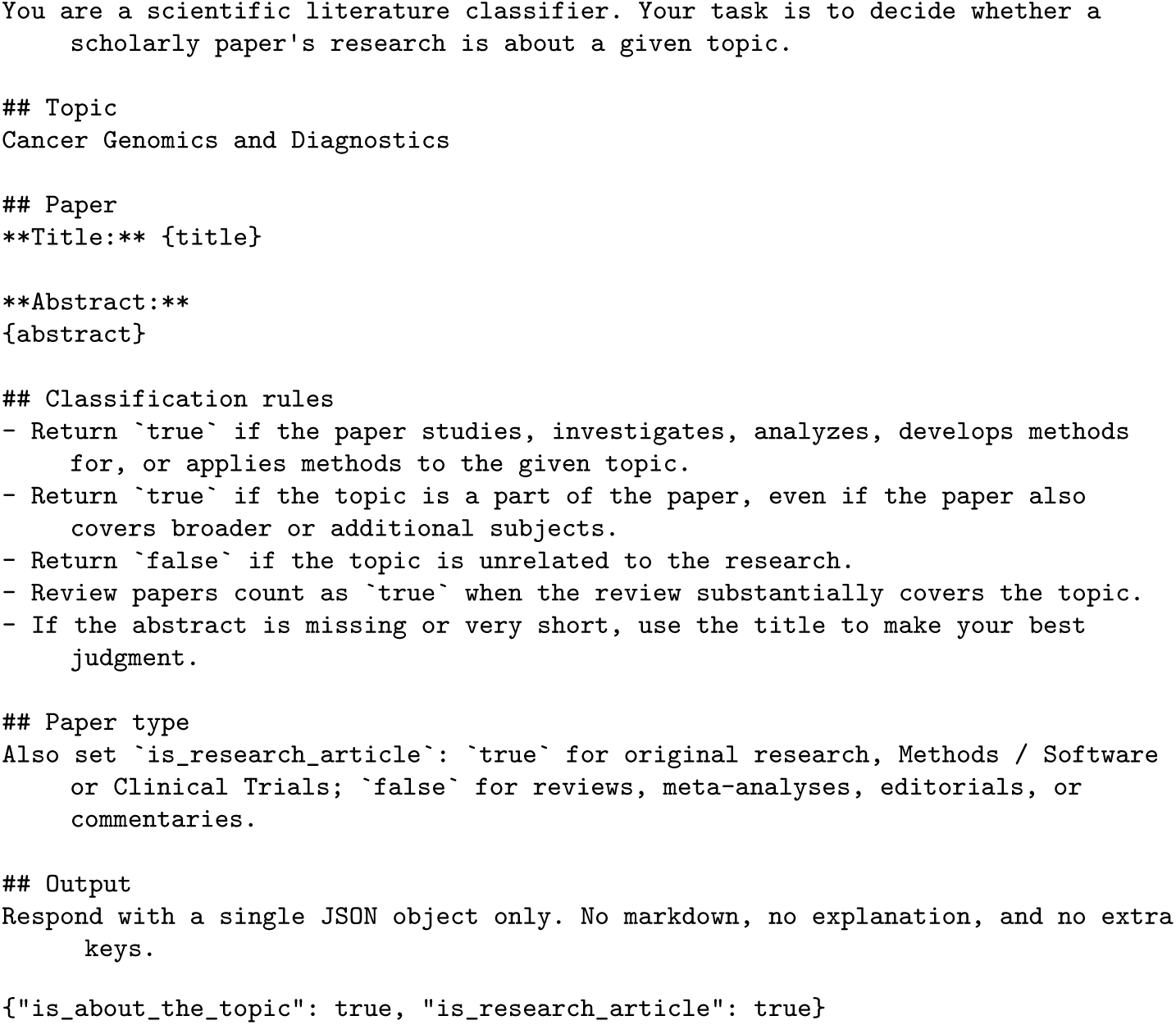

The model was additionally instructed via a system prompt to respond with a single valid JSON object containing only the keys is_about_the_topic and is_research_article, each with a boolean value. Papers classified as on-topic research articles were retained; those classified as off-topic or as non-research document types were excluded. Classification was performed using Claude Sonnet 5 (Anthropic), accessed via API. For each cohort, the 90 highest-ranked candidates by 18-month citation count were evaluated. Papers failing either classification criterion were excluded, and the top 50 on-topic papers by 18-month citation count were retained for downstream analysis. In the Class of 2025 cohort, 68 of 90 evaluated papers passed the relevance filter and 22 were excluded; in the Class of 2026 cohort, 61 of 90 passed and 29 were excluded. In both cohorts the initial 1.8× candidate pool multiplier was sufficient to yield 50 on-topic papers without requiring expansion of the pool multiplier.

### Concept extraction

Scientific concepts were extracted from the titles and abstracts of all retained papers using an LLM-based extraction pipeline^22^. For each paper, a structured prompt instructed the model to identify scientifically important concepts central to the paper’s content, spanning disease entities, molecular targets, genes and biomarkers, biological mechanisms and pathways, cell types, therapeutic strategies, experimental and computational methods, data types, and emerging research directions. The model was instructed to preserve the original terminology used by the authors, to prefer specific over generic terms, and not to normalize synonyms at this stage. Up to 20 concepts were requested per paper. Concept extraction was performed using Claude Sonnet 5 (Anthropic), accessed via API, with responses parsed as JSON arrays of concept name strings. Concepts were collected across all 100 papers in the combined corpus, yielding 1,364 unique raw concept strings after case-insensitive deduplication before normalization.

### Concept normalization and canonical taxonomy construction

Raw extracted concepts were normalized into a shared two-level taxonomy through an LLM pipeline^23^. In the first stage (canonical mapping), all case-insensitively deduplicated concept strings were grouped into synonym clusters representing the same or closely related scientific ideas. To ensure that semantically related terms were preferentially grouped within the same LLM batch, concept strings were first embedded using Gemini Embedding 001 (Google)^24^, and similarity-based clustering was performed by computing pairwise cosine similarity between term embeddings and grouping terms into connected components using a cosine similarity threshold of 0.78. Similarity-connected components were packed into batches of 200–400 terms. Each batch was submitted to DeepSeek V4 Pro (DeepSeek), accessed via API, with instructions to identify synonym groups and assign a canonical concept name—the most commonly recognized scientific term—to each group; singleton terms with no identified synonyms were retained under their own name. This single-round canonical mapping procedure produced 1,090 canonical concept groups from 1,364 deduplicated raw concept strings across the combined corpus.

### Parent theme assignment

In the second normalization stage (parent mapping), each canonical concept name was assigned to a higher-level scientific parent theme^25^. Canonical names were again embedded and clustered by cosine similarity using the same similarity threshold and batch size parameters as the canonical mapping stage, ensuring that related canonical concepts were processed within the same LLM batch. Each batch was submitted to DeepSeek V4 Pro with instructions to assign every canonical name to exactly one broader parent category representing a meaningful scientific theme. Any canonical names not assigned by the initial pass were submitted in a follow-up batch. This stage produced 77 parent themes with complete coverage across all 1,090 canonical concept groups.

### Concept frequency comparison across cohorts

For each paper in the normalized corpus, all extracted concepts were mapped to their canonical name and parent theme through the full two-level taxonomy. Concept occurrence counts were tabulated separately for the Class of 2025 (2023 publications) and Class of 2026 (2024 publications) cohorts at both the canonical-topic and parent-theme levels. For each paper, concept occurrence was counted at most once per canonical topic or parent theme (within-paper deduplication). Raw counts represent the number of papers in each cohort in which a given concept or theme appeared. Counts were normalized by the total number of papers in each cohort (50 for Class of 2025; 50 for Class of 2026), yielding paper-level prevalence (proportion of papers containing each concept or theme). The normalized frequency difference (Class of 2026 minus Class of 2025), fold change of normalized frequencies, and log_2 fold change were computed for each concept and theme to quantify shifts in thematic representation between cohorts. A small constant (1 × 10 ¹²) was added to both normalized frequencies when computing fold change to avoid division by zero. Trend visualizations highlighted the ten concepts or themes with the largest positive and negative frequency differences in diverging bar charts and scatter plots, and the fifteen concepts with the highest Class of 2026 normalized frequency in grouped bar charts, separately at the canonical-topic and parent-theme levels. No statistical significance threshold was applied to the frequency comparisons, which are descriptive in nature; reported values are paper-level prevalences and their differences.

### Word cloud visualization

To characterize the thematic content of the most highly cited papers in the Class of 2026 cohort, word clouds^26^ were generated from the concept frequency data of the 50 highest-ranked papers by 18-month citation count. Word size in each cloud was scaled proportionally to the number of papers in the selected set mentioning each canonical topic or parent theme, with each concept counted at most once per paper, separately at the canonical-topic and parent-theme levels.

### Software and reproducibility

All analyses were performed in Python 3.12.13 using pandas 3.0.3^27^, NumPy 2.4.4^28^, matplotlib 3.11.0^29^, and WordCloud 1.9.6. Figures were generated with publication settings (Arial font, 300 dpi for raster outputs, SVG for vector outputs). Bibliographic queries were executed against the OpenAlex REST API in July 2026. LLM tasks used Claude Sonnet 5 for topical screening and concept extraction, DeepSeek V4 Pro for canonical and parent-theme normalization, and Gemini Embedding 001 for similarity-based batching. Complete ranked paper lists, extracted and normalized concepts, frequency tables, and figure-generation inputs are archived for independent reproduction (Zenodo DOI: 10.5281/zenodo.21436170).

## Results

### Literature corpus and cohort assembly

Bibliographic retrieval from OpenAlex^16^ under topic T11287 (Cancer Genomics and Diagnostics) identified 10,175 research articles published between 1 January and 31 December 2024, and 11,081 research articles published in 2023, that met the inclusion criteria. After ranking by 18-month citation count and large language model (LLM)–based topical relevance filtering, 50 on-topic articles were retained from each cohort, yielding a combined analytic corpus of 100 papers. For the Class of 2026 cohort (2024 publications), 90 candidate papers were evaluated for topical relevance; 61 were classified as on-topic and 29 were excluded. For the Class of 2025 cohort (2023 publications), 90 papers were evaluated, of which 68 passed the relevance filter and 22 were excluded. In both cohorts, the 1.8× LLM pool multiplier was sufficient to yield at least 50 on-topic papers without expansion.

### Early citation impact of top-ranked papers

Among the 50 highest-ranked on-topic papers in each cohort, 18-month citation counts ranged from 50 to 340 in the Class of 2025 cohort and from 51 to 412 in the Class of 2026 cohort, with a median of 76 citations in the Class of 2025 cohort and 73.5 citations in the Class of 2026 cohort. The most cited paper in the Class of 2025 cohort was “Analysis and Visualization of Longitudinal Genomic and Clinical Data from the AACR Project GENIE Biopharma Collaborative in cBioPortal” (340 citations; *Cancer Research*)^30^. The most cited paper in the Class of 2026 cohort was “Genomic data in the All of Us Research Program” (412 citations; *Nature*)^31^.

High-impact journals were strongly represented in both cohorts. In the Class of 2025 cohort, 30 of 50 papers were published in Nature portfolio journals, including 13 in *Nature*, 8 in *Nature Medicine*, 4 in *Nature Communications*, 3 in *Nature Genetics*, 1 in *Nature Machine Intelligence*, and 1 in *Nature Biotechnology*. In the Class of 2026 cohort, 31 of 50 papers appeared in Nature portfolio journals, including 13 in *Nature*, 6 in *Nature Medicine*, 4 in *Nature Cancer*, 3 in *Nature Communications*, 2 in *Nature Genetics*, 1 in *Nature Protocols*, 1 in *Nature Machine Intelligence*, and 1 in *Nature Biotechnology*. Other frequently represented journals included *Cancer Cell* (3 papers in each cohort) and *Annals of Oncology* (1 paper in each cohort). The complete ranked lists of the top 50 on-topic papers for each cohort are provided in Supplementary Tables S1^32–81^ and S2^31,82–130^.

### Concept extraction and taxonomy construction

LLM-based concept extraction from the titles and abstracts of all 100 retained papers yielded 1,364 unique raw concept strings after case-insensitive deduplication. Normalization into a shared two-level taxonomy produced 1,090 canonical concept groups and 77 parent themes, with complete mapping coverage across the corpus. The Class of 2025 cohort contributed 817 concept mentions across 50 papers (mean, 16.3 per paper), and the Class of 2026 cohort contributed 803 concept mentions across 50 papers (mean, 16.1 per paper).

### Thematic landscape of Class of 2026 top-cited papers

Word clouds generated from the 50 highest-ranked Class of 2026 papers by 18-month citation count characterize the dominant scientific themes of the most influential recent publications at the parent-theme level (Figure 1) and canonical-topic level (Figure 2). At the parent-theme level, the most frequently represented categories were Cancer types and histopathological classification (31 papers), Clonal evolution and intratumoral heterogeneity (19 papers), and Tumor microenvironment and cancer-associated fibroblasts (12 papers). At the canonical-topic level, tumor heterogeneity was the single most frequent concept (10 papers), followed by tumor microenvironment (9 papers), and immune cell infiltration, copy number variation, and single-cell RNA sequencing (7 papers each).

**Figure 1.**
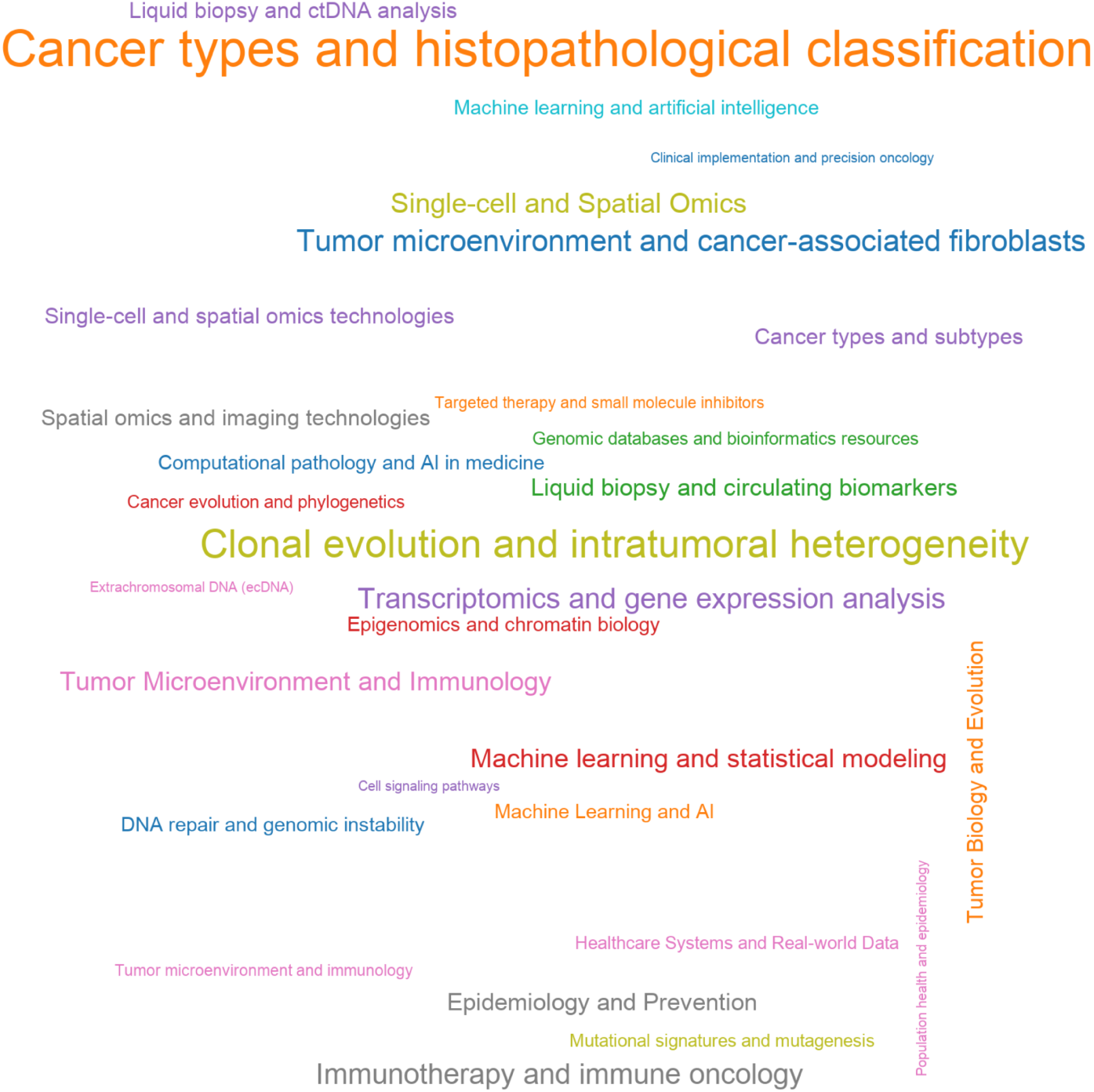
Word cloud of parent themes across the 50 most-cited Class of 2026 papers. Word size is proportional to the number of papers containing each parent theme among the 50 papers with the highest 18-month citation counts in the Class of 2026 cohort. Labels denote parent-theme categories assigned during two-level concept normalization. Class of 2026, publication cohort comprising articles published in 2024.

**Figure 2.**
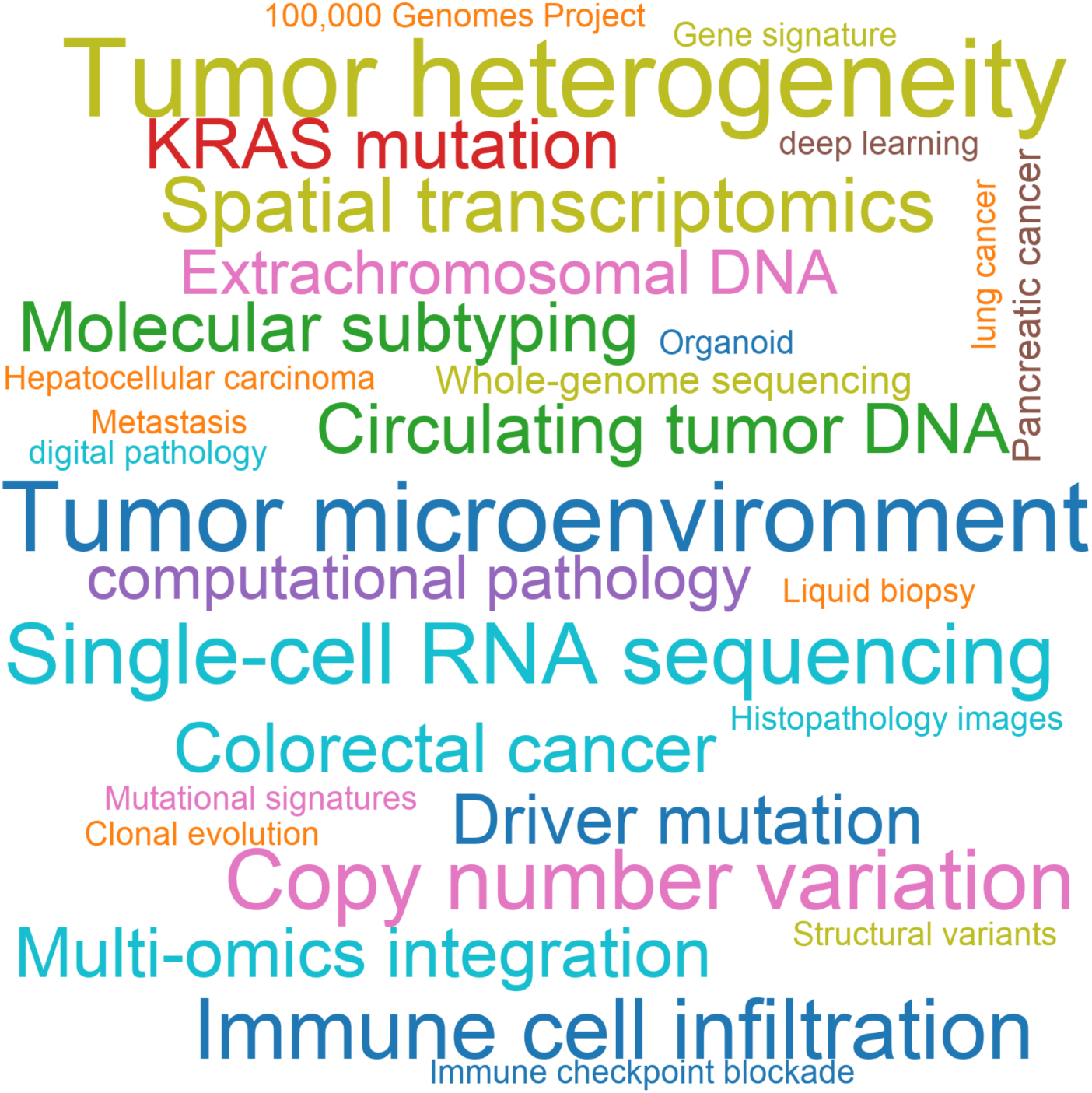
Word cloud of canonical topics across the 50 most-cited Class of 2026 papers. Word size is proportional to the number of papers containing each canonical topic among the 50 papers with the highest 18-month citation counts in the Class of 2026 cohort. Labels denote canonical concept names assigned during two-level concept normalization. Class of 2026, publication cohort comprising articles published in 2024.

### Cohort comparison at the parent-theme level

Comparison of normalized concept frequencies between the Class of 2025 and Class of 2026 cohorts revealed systematic shifts in the thematic composition of highly cited publications (Figure 3). The diverging bar chart (Figure 3a) highlighted parent themes with the largest cohort differences. Among themes with the largest increases in normalized frequency in the Class of 2026 cohort, Clonal evolution and intratumoral heterogeneity showed the greatest gain (frequency difference, +0.140; papers containing the theme, 12 in Class of 2025 versus 19 in Class of 2026), followed by Single-cell and spatial omics technologies (+0.100; 2 versus 7), Spatial omics and imaging technologies (+0.080; 3 versus 7), Tumor microenvironment and cancer-associated fibroblasts (+0.060; 9 versus 12), and Epidemiology and Prevention (+0.060; 5 versus 8).

**Figure 3.**
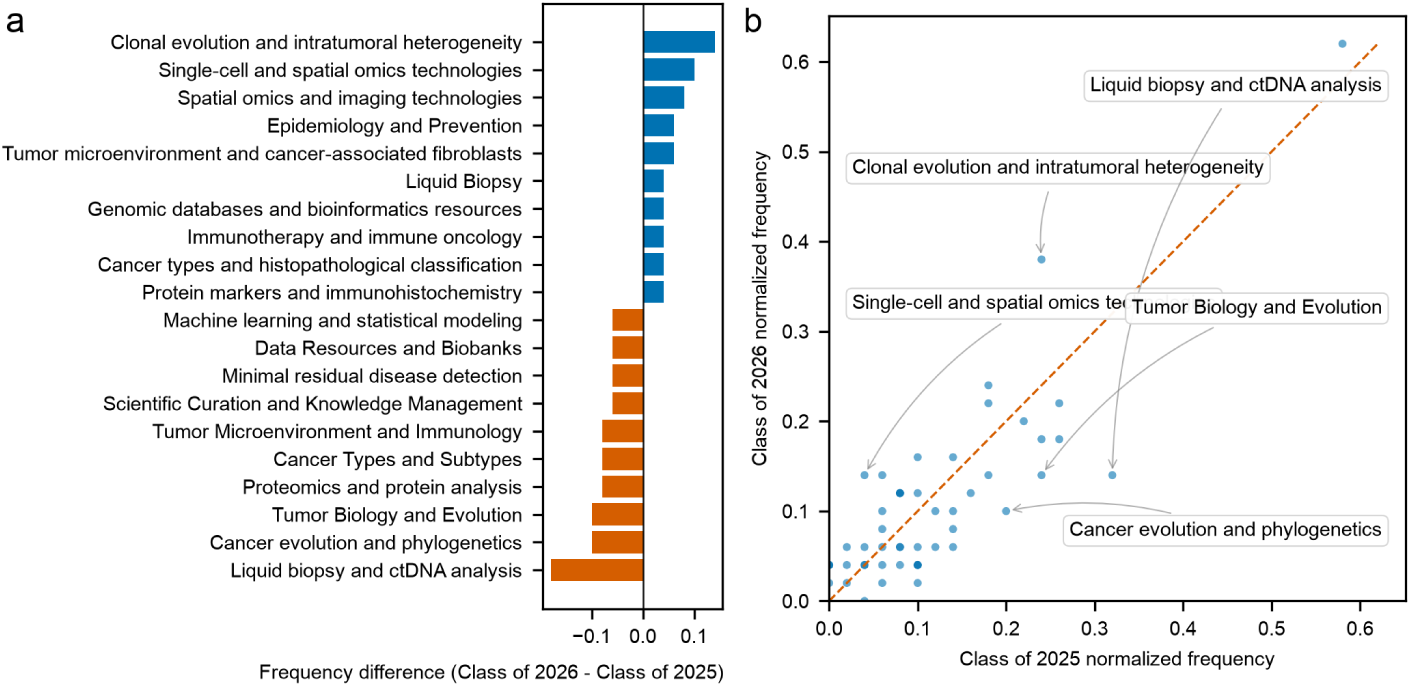
Cohort comparison of concept frequencies at the parent-theme level. (3a) Diverging bar chart showing the 10 parent themes with the largest positive and 10 with the largest negative normalized frequency differences between cohorts. Bars extend rightward (blue) for themes more frequent in the Class of 2026 cohort and leftward (orange) for themes more frequent in the Class of 2025 cohort; the x-axis represents the normalized frequency difference (Class of 2026 minus Class of 2025). (3b) Scatter plot in which each point represents one parent theme; the x-axis shows normalized frequency in the Class of 2025 cohort and the y-axis shows normalized frequency in the Class of 2026 cohort. The dashed diagonal line indicates equal normalized frequency in both cohorts; points above the line are more frequent in the Class of 2026 cohort and those below are more frequent in the Class of 2025 cohort. The 10 themes with the largest absolute frequency differences are annotated. Class of 2025, publication cohort comprising articles published in 2023; Class of 2026, publication cohort comprising articles published in 2024.

Among parent themes with the largest decreases (Figure 3a), Liquid biopsy and ctDNA analysis declined most markedly (frequency difference, -0.180; 16 versus 7), followed by Cancer evolution and phylogenetics (-0.100; 10 versus 5), Tumor Biology and Evolution (-0.100; 12 versus 7), Proteomics and protein analysis (-0.080; 5 versus 1), Cancer Types and Subtypes (-0.080; 7 versus 3), and Tumor Microenvironment and Immunology (-0.080; 13 versus 9). The scatter plot (Figure 3b) showed that most parent themes clustered near the diagonal of equal normalized frequency, with annotated outliers corresponding to the themes with the largest absolute frequency differences described above.

Among the 15 parent themes with the highest normalized frequency in the Class of 2026 cohort (Figure 4), Cancer types and histopathological classification and Clonal evolution and intratumoral heterogeneity ranked highest (normalized frequency, 0.620 and 0.380, respectively), followed by Tumor microenvironment and cancer-associated fibroblasts (0.240), Immunotherapy and immune oncology (0.220), Transcriptomics and gene expression analysis (0.220), and Single-cell and Spatial Omics (0.200). Of these leading themes, Cancer types and histopathological classification, Clonal evolution and intratumoral heterogeneity, Tumor microenvironment and cancer-associated fibroblasts, Immunotherapy and immune oncology, Epidemiology and Prevention, Liquid biopsy and circulating biomarkers, Single-cell and spatial omics technologies, and Spatial omics and imaging technologies showed higher normalized frequency in the Class of 2026 cohort than in the Class of 2025 cohort; Transcriptomics and gene expression analysis, Single-cell and Spatial Omics, Machine learning and statistical modeling, Tumor Microenvironment and Immunology, Cancer types and subtypes, Tumor Biology and Evolution, and Liquid biopsy and ctDNA analysis were more represented in the Class of 2025 cohort.

**Figure 4.**
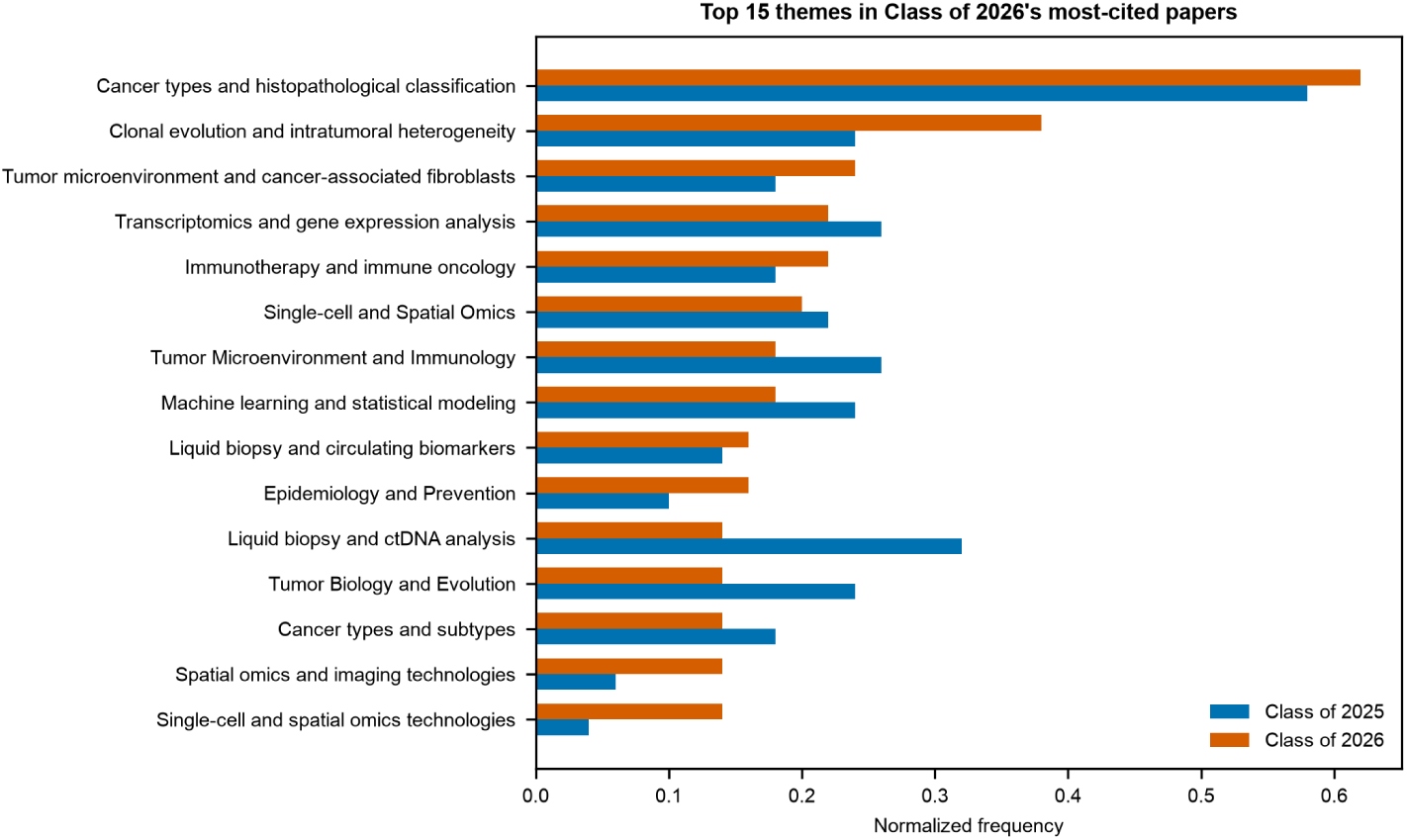
Normalized frequencies of the 15 most prevalent parent themes in the Class of 2026 cohort. Grouped horizontal bar chart showing normalized concept frequency for the 15 parent themes with the highest normalized frequency in the Class of 2026 cohort. Blue bars indicate the Class of 2025 cohort; orange bars indicate the Class of 2026 cohort. Themes are ordered by descending normalized frequency in the Class of 2026 cohort. Normalized frequency is the number of papers containing a given theme divided by 50 (paper-level prevalence). Class of 2025, publication cohort comprising articles published in 2023; Class of 2026, publication cohort comprising articles published in 2024.

The complete parent-theme frequency comparison, including paper counts, normalized frequencies, frequency differences, fold changes, and log_2 fold changes for all 77 parent themes, is provided (Supplementary Table S3).

### Cohort comparison at the canonical-topic level

At the canonical-topic level, comparison of normalized concept frequencies between cohorts revealed similar thematic shifts (Figure 5). The diverging bar chart (Figure 5a) showed that copy number variation and immune cell infiltration exhibited the largest increases in normalized frequency (frequency difference, +0.100 each; papers containing the topic, 2 in Class of 2025 versus 7 in Class of 2026). Spatial transcriptomics showed the next-largest gain (+0.100; 1 versus 6), followed by driver mutation (+0.080; 1 versus 5), KRAS mutation (+0.060; 2 versus 5), colorectal cancer (+0.060; 2 versus 5), and the 100,000 Genomes Project (+0.060; 0 versus 3).

**Figure 5.**
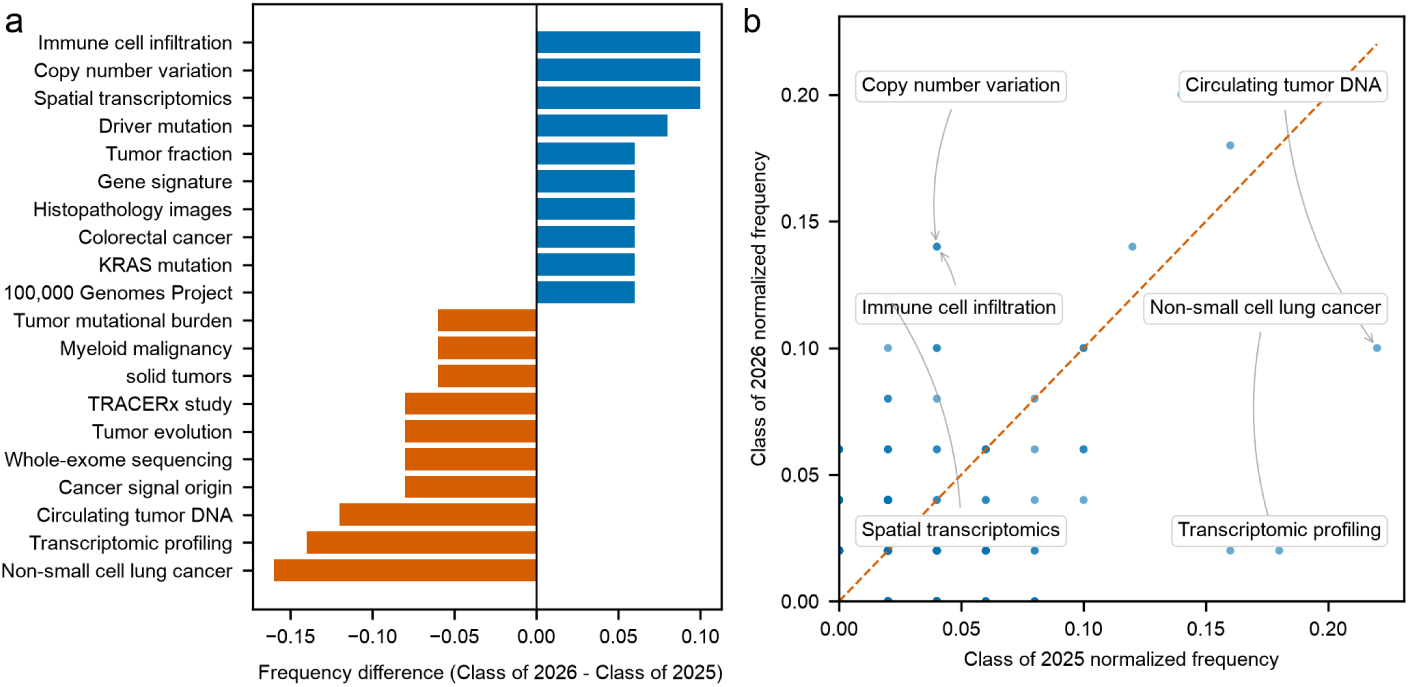
Cohort comparison of concept frequencies at the canonical-topic level. (5a) Diverging bar chart showing the 10 canonical topics with the largest positive and 10 with the largest negative normalized frequency differences between cohorts. Bars extend rightward (blue) for topics more frequent in the Class of 2026 cohort and leftward (orange) for topics more frequent in the Class of 2025 cohort; the x-axis represents the normalized frequency difference (Class of 2026 minus Class of 2025). (5b) Scatter plot in which each point represents one canonical topic; the x-axis shows normalized frequency in the Class of 2025 cohort and the y-axis shows normalized frequency in the Class of 2026 cohort. The dashed diagonal line indicates equal normalized frequency; points above the line are more frequent in the Class of 2026 cohort and those below are more frequent in the Class of 2025 cohort. The 10 topics with the largest absolute frequency differences are annotated. Class of 2025, publication cohort comprising articles published in 2023; Class of 2026, publication cohort comprising articles published in 2024.

Canonical topics with the largest decreases (Figure 5a) included non-small cell lung cancer (frequency difference, -0.160; 9 versus 1), transcriptomic profiling (-0.140; 8 versus 1), circulating tumor DNA (-0.120; 11 versus 5), cancer signal origin (-0.080; 4 versus 0), whole-exome sequencing (-0.080; 4 versus 0), and tumor evolution (-0.080; 4 versus 0). As in the parent-theme analysis, the scatter plot (Figure 5b) placed most canonical topics near the diagonal, with annotated outliers marking the topics with the largest absolute frequency differences.

Among the 15 canonical topics with the highest normalized frequency in the Class of 2026 cohort (Figure 6), tumor heterogeneity ranked highest (0.200), followed by tumor microenvironment (0.180), copy number variation (0.140), immune cell infiltration (0.140), single-cell RNA sequencing (0.140), and spatial transcriptomics (0.120).

**Figure 6.**
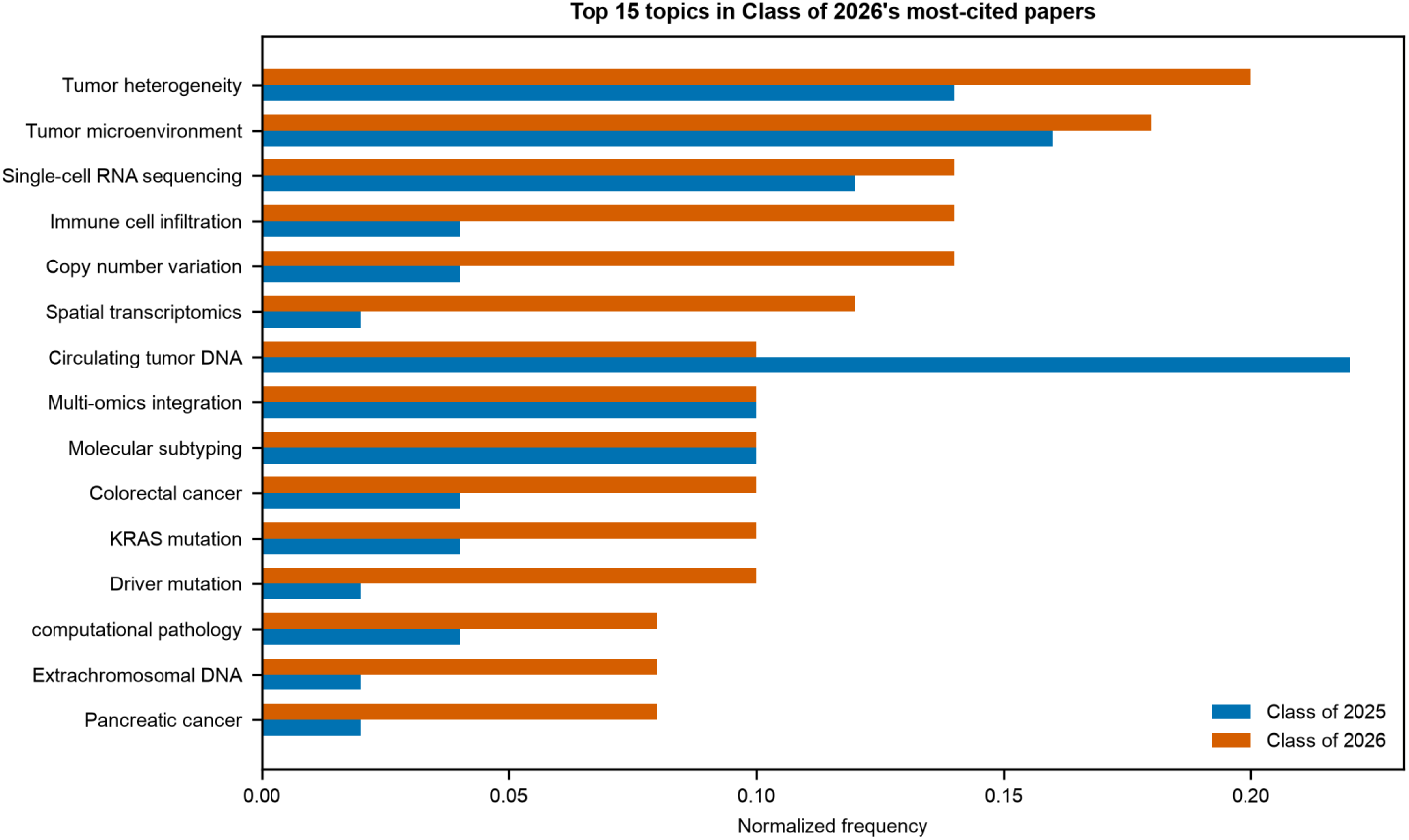
Normalized frequencies of the 15 most prevalent canonical topics in the Class of 2026 cohort. Grouped horizontal bar chart showing normalized concept frequency for the 15 canonical topics with the highest normalized frequency in the Class of 2026 cohort. Blue bars indicate the Class of 2025 cohort; orange bars indicate the Class of 2026 cohort. Topics are ordered by descending normalized frequency in the Class of 2026 cohort. Normalized frequency is the number of papers containing a given topic divided by 50 (paper-level prevalence). Class of 2025, publication cohort comprising articles published in 2023; Class of 2026, publication cohort comprising articles published in 2024.

The complete canonical-topic frequency comparison for all 1,090 canonical concept groups is provided (Supplementary Table S4).

## Discussion

By integrating 18-month citation ranking, large language model (LLM)–based topical relevance filtering, and a two-level concept taxonomy, this study mapped the scientific landscape of the most influential recent publications in Cancer Genomics and Diagnostics across two consecutive cohorts. The principal finding is twofold. First, early citation impact among top-ranked papers is disproportionately driven by large-scale genomic infrastructure and data-access platforms rather than by single-disease mechanistic studies alone. Second, the thematic composition of these influential papers is shifting: compared with the Class of 2025 cohort (2023 publications), the Class of 2026 cohort (2024 publications) shows greater representation of clonal evolution and intratumoral heterogeneity, single-cell and spatial omics technologies, spatial transcriptomics, immune cell infiltration, and copy number variation, alongside persistent emphasis on cancer types and histopathological classification, tumor heterogeneity, and tumor microenvironment biology. These patterns suggest that the field is consolidating around integrative, spatially resolved, and heterogeneity-aware approaches to genomic medicine while retaining core preoccupations with histologic stratification and microenvironmental context.

The dominance of infrastructure-oriented papers among the highest early-cited works is itself informative. The most cited paper in the Class of 2026 cohort describes the release of clinical-grade whole-genome sequences from the All of Us Research Program, linked to longitudinal electronic health records and characterized by substantial ancestral diversity^31^. The leading paper in the Class of 2025 cohort reports enhanced visualization and longitudinal analysis capabilities for AACR Project GENIE clinico-genomic data within cBioPortal^30^, building on GENIE’s role as one of the largest publicly available real-world oncology genomics registries^9^. That both cohort leaders are resource and tooling papers—not landmark therapeutic trials—indicates that the research community assigns substantial early citation weight to reusable datasets and analytical platforms that enable downstream discovery across many laboratories and disease contexts. The strong representation of Nature portfolio journals in both cohorts further reflects the visibility accorded to such foundational releases, although this concentration also cautions against equating journal prestige with field-wide representativeness.

Several themes that gained prevalence in the Class of 2026 cohort align with broader trajectories in precision oncology. At the parent-theme level, the largest increase was observed for clonal evolution and intratumoral heterogeneity, consistent with growing recognition that subclonal architecture and intratumoral diversity are central to prognosis, therapeutic resistance, and biomarker interpretation^39,48^. Gains in single-cell and spatial omics technologies and spatial omics and imaging technologies mirror accelerating adoption of spatial transcriptomics for resolving tumor architecture in situ^43,94^. At the canonical level, spatial transcriptomics, immune cell infiltration, and copy number variation showed among the largest positive frequency differences, reinforcing a shift toward diagnostics that integrate spatial immune context with structural genomic alteration profiling^7,131^. Tumor heterogeneity and tumor microenvironment ranked highest among canonical topics in the Class of 2026 word cloud, while cancer types and histopathological classification remained the dominant parent theme, underscoring that histologic stratification remains foundational even as spatial and heterogeneity-oriented modalities expand.

Conversely, several themes were more prevalent among Class of 2025 than Class of 2026 top-cited papers, and these declines warrant careful interpretation. Liquid biopsy and ctDNA analysis showed the largest negative frequency difference at the parent-theme level, and circulating tumor DNA declined at the canonical level. This pattern should not be read as evidence that liquid biopsy is losing clinical relevance; comprehensive reviews continue to emphasize ctDNA as a minimally invasive modality for mutation detection, treatment monitoring, and residual disease assessment^132,133^. Rather, the shift likely reflects cohort-specific composition among early-high-impact papers: the Class of 2025 set may have contained a cluster of ctDNA- and MRD-focused trial and methods publications that accumulated citations rapidly within the 18-month window, whereas the Class of 2026 leaderboard was comparatively enriched for population-scale sequencing resources, spatial omics studies, and heterogeneity-oriented integrative analyses. Parallel declines in non-small cell lung cancer and transcriptomic profiling at the canonical level, alongside reductions in cancer evolution and phylogenetics and tumor biology and evolution at the parent-theme level, may similarly reflect year-specific paper mixtures within the top-fifty constraint rather than diminished research activity in lung cancer genomics or evolutionary oncology, for which bibliometric analyses document sustained growth in omics, tumor microenvironment, and immunotherapy-oriented investigations^11^.

Comparison with prior bibliometric studies highlights both convergence and methodological distinction. Analyses of lung cancer omics literature and of AI-integrated multi-omics in cancer research similarly identify tumor microenvironment profiling and machine learning as dominant or accelerating themes^11,13^. Our findings are broadly concordant with these signals while additionally highlighting spatial transcriptomics, immune infiltration metrics, and clonal heterogeneity among the most early-cited on-topic work. Most prior studies employ keyword co-occurrence or citation network analyses over large corpora, whereas the present framework ranks papers by early citation impact, filters for topical relevance, and extracts author-preserving concepts through an embedding-guided LLM taxonomy. This design prioritizes near-term influence among on-topic work and may detect semantic shifts obscured when all publications are weighted equally.

This study contributes a reproducible, semantically enriched bibliometric framework with several distinctive features. The threshold-expansion algorithm for 18-month citation ranking provides a mathematically exact top-N leaderboard without exhaustive scoring of the full OpenAlex corpus. LLM-based relevance filtering addresses known imprecision in indexer-assigned topic labels, and the two-stage concept normalization pipeline yields a structured taxonomy amenable to cohort comparison. Using OpenAlex as an open bibliographic source further supports transparency and replicability in line with growing adoption of open scholarly indexes in bibliometric research^14^. At the same time, the application of LLMs for classification and concept extraction introduces considerations familiar from scientometric applications of generative models: although LLMs can scale annotation tasks efficiently, their outputs benefit from external validation and should be interpreted as structured approximations rather than ground truth^134^.

The translational implications of the observed thematic shifts are substantial. Diagnostic laboratories may need to expand beyond panel-based sequencing toward workflows integrating spatial immune profiling, copy-number interpretation, and single-cell or spatial transcriptomic readouts with routine molecular results^4,7^. For research funders, the citation prominence of diversity-enriched, EHR-linked genomic resources underscores the multiplier effect of open, well-annotated cohorts^31^. Clinicians and translational researchers should note the co-emergence of clonal heterogeneity, spatial omics, and immune-context biomarkers as indicators of continued emphasis on resolving tumor complexity for therapeutic decision-making^135^.

Several strengths and limitations frame the interpretation of these findings. Strengths include completed 18-month citation windows for both cohorts, exact ranking methodology, LLM topical filtering with reported exclusion rates, and two-level concept mapping across one hundred retained papers. Limitations include descriptive frequency comparisons without formal statistical testing, citation counts as proxies for visibility rather than quality, and potential early-citation advantage for infrastructure papers. Concept extraction from titles and abstracts alone cannot capture full-text detail, and LLM-based normalization may introduce model-dependent synonym groupings not validated by expert curators. Finally, the analytic corpus comprises only the top fifty on-topic papers per year from a topic classification encompassing more than ten thousand annual research articles; generalization to the broader literature therefore requires caution^14^.

Future work should extend this framework across additional publication cohorts and longer citation horizons to distinguish transient attention from durable influence. Expert validation of the LLM-derived taxonomy, incorporation of full-text and citation-context analysis, and linkage to clinical trial registries and funding databases would strengthen causal interpretation of thematic change. Statistical testing of theme prevalence differences, longitudinal tracking of spatial omics–liquid biopsy convergence, and periodic updating of the Publication Rank Index would transform this descriptive snapshot into a longitudinal monitoring system for Cancer Genomics and Diagnostics^136^.

In summary, the most influential recent publications in Cancer Genomics and Diagnostics are increasingly defined by population-scale genomic infrastructure, spatially resolved and single-cell omics profiling, clonal heterogeneity science, and immune-context biomarker integration, even as core concerns with histopathological classification, tumor microenvironment biology, and copy-number alteration profiling persist. The observed retreat of liquid-biopsy-centric themes from the top-cited set between consecutive cohorts reflects shifting citation composition among high-impact papers rather than waning clinical importance of ctDNA. By coupling exact early-impact citation ranking with LLM-based semantic mapping, this study provides a replicable lens on how a rapidly evolving field prioritizes and integrates genomic discovery, diagnostic innovation, and translational infrastructure—and offers a template for monitoring those priorities as precision oncology continues to mature.

## Acknowledgments

We thank OpenAlex for providing open bibliographic metadata and citation data. LLM-based classification and concept extraction used models from Anthropic and DeepSeek accessed via the OpenRouter API. Analyses and figures were produced with open-source Python tools including pandas and matplotlib.

## Funding

This work was self-funded; no external funding was received.

## Data availability

The ranked paper lists, concept frequencies, and normalization outputs underlying this analysis are available on Zenodo at https://doi.org/10.5281/zenodo.21436170. Source code and algorithm documentation are available at https://github.com/pepkio/pri-top-papers. A web interface for exploring the results is available at https://pri.pepkio.com/.

## Competing interests

The authors declare no competing interests.

## Supplementary Tables

**Supplementary Table S1.** Ranked list of the 50 highest 18-month-citation on-topic papers in the Class of 2025 cohort (2023 publications). Columns report rank, 18-month citation count, topical relevance classification, article title, journal, publication date, and DOI.

**Supplementary Table S2.** Ranked list of the 50 highest 18-month-citation on-topic papers in the Class of 2026 cohort (2024 publications). Columns report rank, 18-month citation count, topical relevance classification, article title, journal, publication date, and DOI.

**Supplementary Table S3.** Parent-theme frequency comparison between the Class of 2025 and Class of 2026 cohorts. Columns report parent-theme name, number of papers containing the theme in each cohort (count_2023, count_2024), normalized frequency in each cohort (norm_freq_2023, norm_freq_2024), normalized frequency difference (freq_diff), fold change (fold_change), and log_2 fold change (log2_fold_change).

**Supplementary Table S4.** Canonical-topic frequency comparison between the Class of 2025 and Class of 2026 cohorts. Columns report canonical topic name, number of papers containing the topic in each cohort (count_2023, count_2024), normalized frequency in each cohort (norm_freq_2023, norm_freq_2024), normalized frequency difference (freq_diff), fold change (fold_change), and log_2 fold change (log2_fold_change).

## Notes

https://doi.org/10.5281/zenodo.21436170

https://github.com/pepkio/pri-top-papers

https://pri.pepkio.com/

### Competing Interest Statement

The authors have declared no competing interest.

### Summary of Updates

Updated the paper filtering and topics filtering criteria and results.

